# Efflux pumps and membrane permeability contribute to intrinsic antibiotic resistance in *Mycobacterium abscessus*

**DOI:** 10.1101/2024.08.23.609441

**Authors:** Kerry McGowen, Tobias Funck, Xin Wang, Samuel Zinga, Ian D. Wolf, Chidiebere C. Akusobi, Claudia M. Denkinger, Eric J. Rubin, Mark R. Sullivan

## Abstract

*Mycobacterium abscessus* is a pulmonary pathogen that exhibits intrinsic resistance to antibiotics, but the factors driving this resistance are incompletely understood. Insufficient intracellular drug accumulation could explain broad-spectrum resistance, but whether antibiotics fail to accumulate in *M. abscessus* and the mechanisms required for drug exclusion remain poorly understood. We measured antibiotic accumulation in *M. abscessus* using mass spectrometry and found a wide range of drug accumulation across clinically relevant antibiotics. Of these compounds, linezolid accumulates the least, suggesting that inadequate uptake impacts its efficacy. We utilized transposon mutagenesis screening to identify genes that cause linezolid resistance and found multiple transporters that promote membrane permeability or efflux, including an uncharacterized, *M. abscessus*-specific protein that effluxes linezolid and several chemically related antibiotics. This demonstrates that membrane permeability and drug efflux are critical mechanisms of antibiotic resistance in *M. abscessus* and suggests that targeting membrane transporters could potentiate the efficacy of certain antibiotics.

## Introduction

*Mycobacterium abscessus* is an opportunistic pathogen that causes lung infection, especially in individuals with structurally abnormal airways as seen with cystic fibrosis, chronic obstructive pulmonary disease, or bronchiectasis^1^. Rates of successful eradication of pulmonary *M. abscessus* infection remain at 30-50% despite aggressive months-long therapy with multiple antibiotics^3,4^. The lack of efficacy of these therapeutic regimens is driven by broad-spectrum antibiotic resistance exhibited by *M. abscessus*^2^, precluding the use of most common antibiotics to treat these infections.

Though *M. abscessus* possesses several mechanisms that confer high-level acquired resistance to specific compounds^3–5^, the extraordinary level of broad antibiotic resistance observed in *M. abscessus* is likely driven by intrinsic mechanisms that reduce drug efficacy. In mycobacteria, one of the most important intrinsic resistance mechanisms is thought to be a highly impermeable, lipid-rich cell wall^6,7^ likely preventing the accumulation of intracellular-acting antibiotics within the cell. Despite the critical nature that this barrier function could play in drug resistance, our understanding of which drugs are most affected by the impermeability of the mycobacterial envelope is incomplete, as the relative accumulation of antibiotics has not been broadly compared in mycobacteria.

Further, the mechanisms by which mycobacteria maintain barrier function against such a broad array of chemicals are incompletely understood. Membrane transporters are thought to play a central role in this process, as they are both important for exporting building blocks of the cell wall and in the direct efflux of drugs from the cell. Numerous membrane transporters have been characterized in mycobacteria, including five distinct superfamilies of transporters: ATP-binding cassette (ABC), major facilitator super family (MFS), small multidrug resistance (SMR), multidrug and toxic-compound extrusion (MATE), and resistance nodulation division (RND) families^8^. Mycobacteria have an abundance of RND-family transporters termed mycobacterial membrane proteins (Mmp) that are typically paired in operons that encode for the large (MmpL) and small (MmpS) units, which together play a critical role in exporting substrates required for the cell envelope^9^. In addition to building the complex mycobacterial cell wall, these families of transporters mobilize a diverse array of substrates and play many physiologically important roles including secretion of virulence factors, adaptation to local environment, and drug efflux^10–14^. While there are multiple examples of mycobacterial proteins that individually promote moderate levels of drug exclusion^10,12,14^, a unified understanding of how the contributions of each of these proteins could result in intrinsic drug resistance remains elusive.

To address these questions, we use mass spectrometry to comparatively analyze which therapeutically relevant antibiotics fail to accumulate in *M. abscessus*, as the potencies of those drugs are more likely to be constrained by ineffective uptake than acquired antibiotic resistance. We then performed a transposon mutagenesis genetic screen with treatment of the antibiotic with the lowest accumulation, linezolid, to examine the mechanisms that drive chemical permeability and intrinsic drug resistance in *M. abscessus*.

## Results

### Therapeutically relevant antibiotics display a wide range of uptake in *M. abscessus*

To assess the relative efficiency of antibiotic accumulation in *M. abscessus*, we developed a liquid chromatography-mass spectrometry (LC-MS) method to simultaneously measure an arrayed panel of 20 antibiotics used to treat mycobacterial infections^15,16^ in the type strain of *M. abscessus subspecies abscessus* (ATC19977) (Fig. 1a). 19 of these antibiotics were detectable over linear ranges that enabled relative quantitation of their uptake in *M. abscessus* (Extended Data Fig. 1a-t). Strikingly, these antibiotics displayed a wide range of intracellular accumulation, with greater than 1000-fold variation between the highest and lowest accumulating compounds (Fig. 1b, Supplementary Table 1). Interestingly, for antibiotics with no known antibiotic resistance mechanisms in *M. abscessus*, there was a negative correlation (R^2^ = 0.62) between intracellular antibiotic accumulation and the average minimum inhibitory concentrations for those drugs (Fig. 1c). This correlation was even stronger (R^2^ = 0.99) when comparing between antibiotics with the same mechanism of action (Fig 1d), suggesting that intracellular accumulation could explain differential efficacy of similarly acting drugs. Together these data suggest that poor intracellular accumulation may be a major determinant of antibiotic efficacy for those compounds.

**Figure 1:**
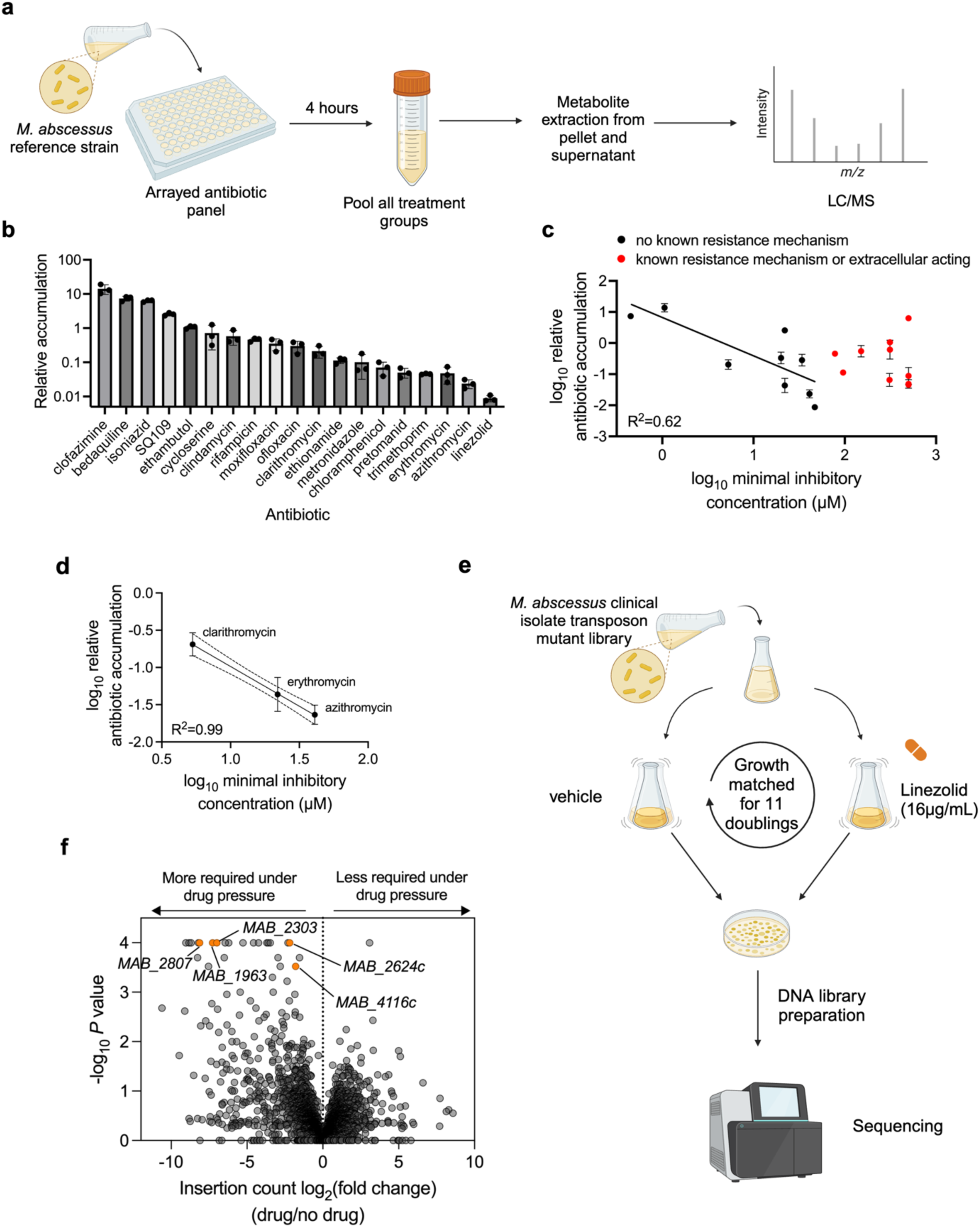
TnSeq reveals differential requirement of numerous membrane transporters with linezolid exposure. **a,** Schematic of antibiotic accumulation assay. **b,** LC-MS measurement of relative accumulation of indicated mycobacterial antibiotics in *M. abscessus* ATC19977 reference strain. Values represent intracellular level of antibiotic after 4 hr incubation normalized to initial antibiotic levels in media prior to incubation. All values are normalized to internal standard and are represented as individual values along with mean ± s.d. n=3 biological replicates. **c,** Minimum inhibitory concentration (MIC50) values of antibiotics with no known acquired resistance mechanisms in *M. abscessus* (black) or with known resistance mechanisms (red) plotted against the relative antibiotic accumulation values as determined in (b). Each individual data point represents a unique antibiotic. R^2^ calculated based on simple linear regression of antibiotics with no known resistance mechanism. **d**, Minimum inhibitory concentration (MIC50) values of macrolides plotted against the relative antibiotic accumulation values as determined in (b). R^2^ calculated based on simple linear regression. **e,** Schematic of TnSeq in *M. abscessus* clinical isolate upon linezolid exposure. **f,** log2-fold ratio of transposon insertion counts plotted against significance with linezolid treatment versus no drug. *P* values derived from two-sided permutation test and are displayed without multiple hypothesis correction. **a**, **e** Created with BioRender.com, released under a Creative Commons Attribution-NonCommercial-NoDerivs 4.0 International license.

We found that linezolid accumulated to a particularly low level (Fig. 1c), suggesting that uptake is a potential hurdle to its efficacy. Linezolid is a bacterial protein synthesis inhibitor that binds to the bacterial 23S ribosomal RNA of the 50S subunit^12^. Linezolid is currently in use as part of an effective regimen for treating multi-drug resistant *Mycobacterium tuberculosis*^17,18^, other opportunistic nontuberculous mycobacteria (NTM) pathogens^16,19^, and many gram-positive pathogens^20^, but it is only moderately effective against *M. abscessus* in liquid culture^21^. However, linezolid has the potential to be an effective co-treatment for *M. abscessus* infections^22,23^, as linezolid synergizes with frontline *M. abscessus* drugs, amikacin and clarithromycin^24,25^, and several studies have demonstrated improved clinical outcomes when linezolid was included in therapeutic regimens^26,27^. Notably, despite multiple inquiries into clinical populations with linezolid resistance, *M. abscessus* fails to show any resistance-associated point mutations in ribosomal linezolid binding sites as have been observed in other bacteria^28^, suggesting that linezolid efficacy could be augmented by improving its intracellular accumulation.

### Linezolid treatment imposes a requirement for numerous membrane transporters

To which identify genes are necessary for *M. abscessus* to survive linezolid and, therefore, likely involved in intrinsic resistance, we performed a genetic screen using transposon mutagenesis and sequencing (TnSeq)^29,30^ (Fig. 1e) in a clinical isolate, *M. abscessus subspecies massiliense* BWH-F^31^ that displays moderate linezolid resistance (Extended Data Fig. 2a). 25 genes were significantly more required (*P* value <0.0005) to survive linezolid treatment compared to the untreated condition (Fig. 1e, Supplemental Table 2-3), and these genes represent a variety of shared functional categories (Table 1)^32^. A large subset of the genes differentially required upon linezolid treatment encode membrane transporters with diverse annotated functions across 3 of the 5 superfamilies of bacterial transporters (Fig. 1f, Extended Data Fig. 2b-f, Table 1). Given the diversity and likely independent functions of these annotated membrane transporter genes, we reasoned that they play a role in efflux or membrane permeability of linezolid and thus impact intrinsic resistance in *M. abscessus*. These genes do not share significant homology across other pathogenic mycobacterial species (Extended Data Table 1)^33^, which suggests the role of most of these genes in linezolid survival is unique to *M. abscessus*. Together, we posited that these membrane proteins each contribute to form an effective, *M. abscessus*-specific permeability barrier or efflux mechanism to limit linezolid accumulation.

**Table 1.**
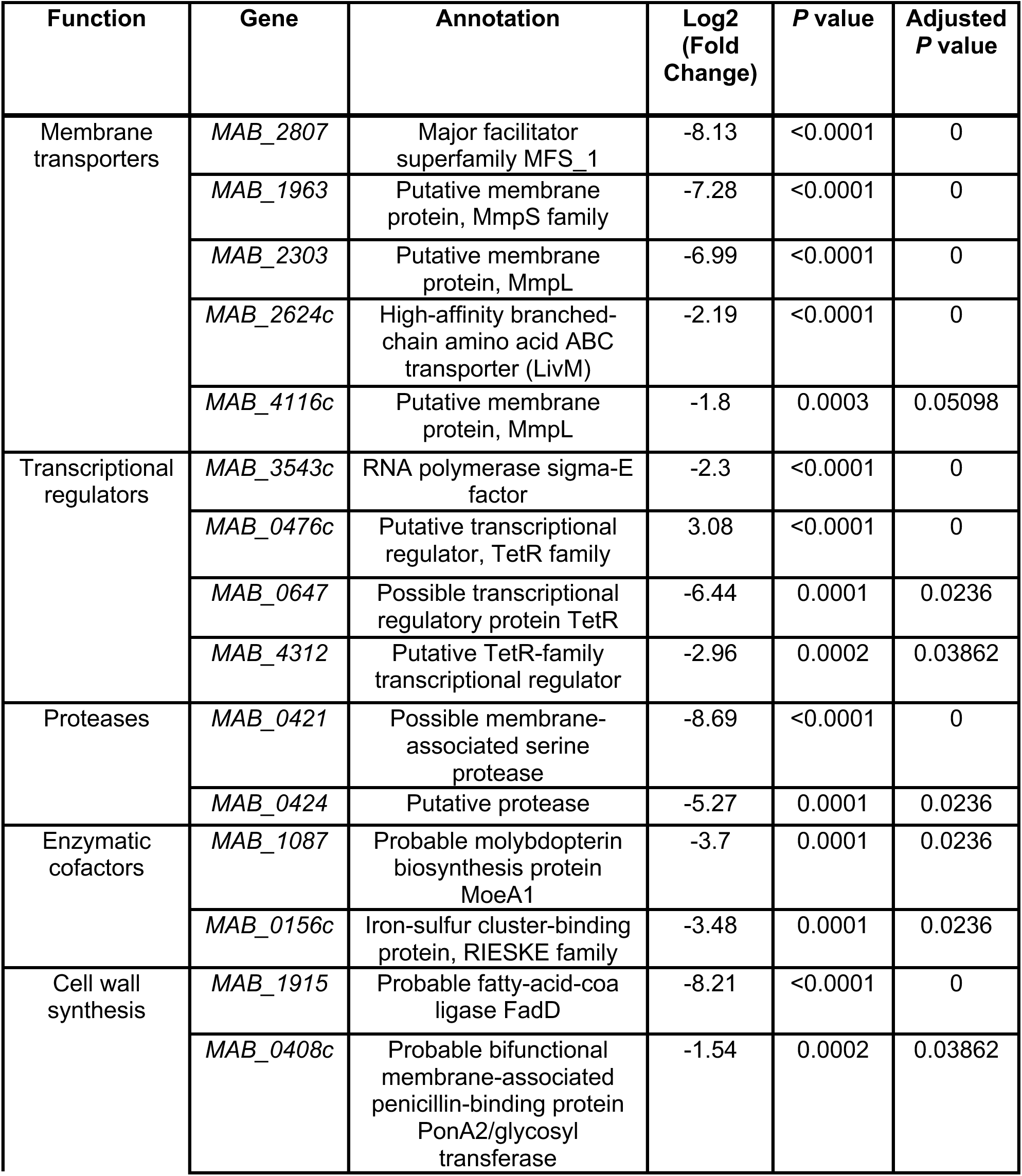
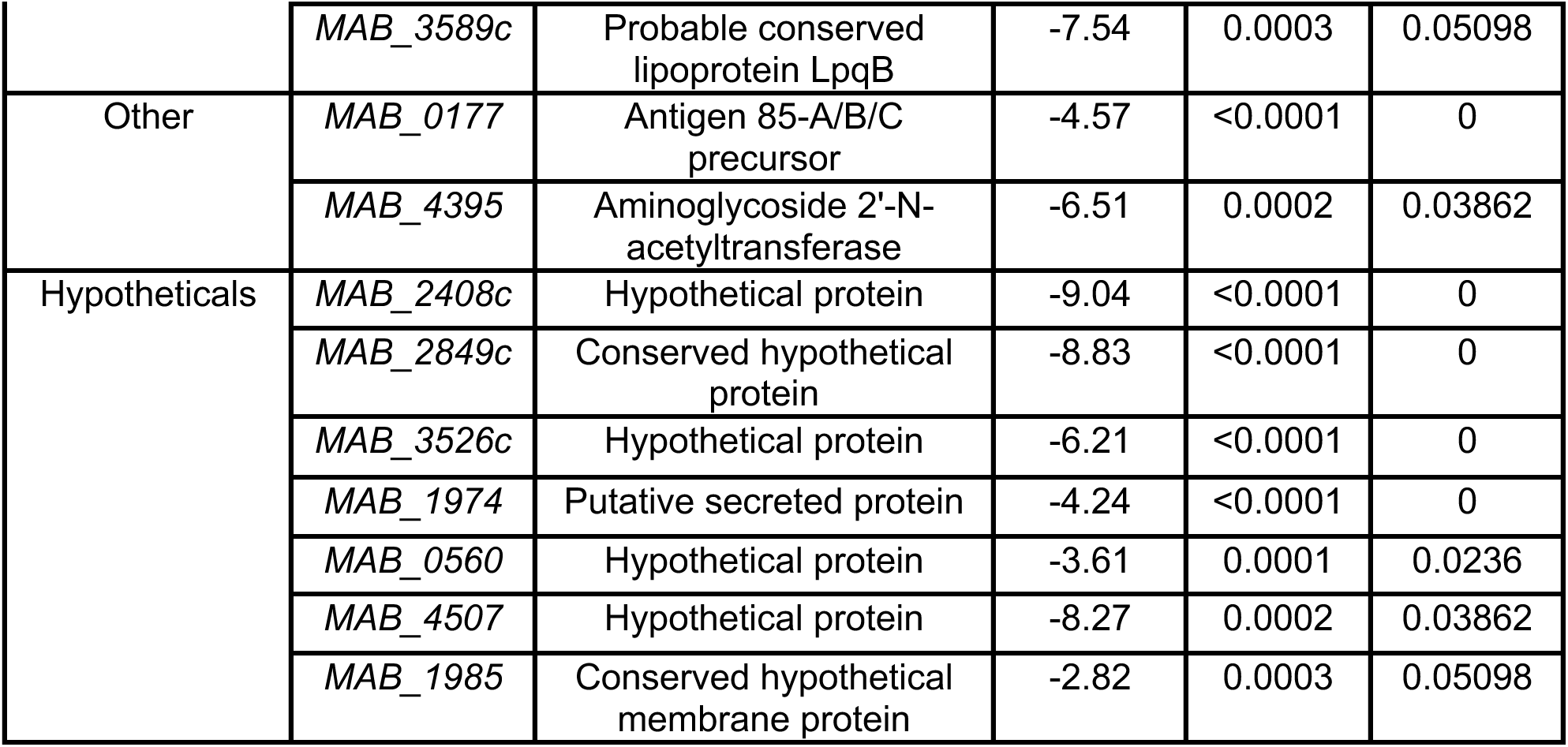
List of genes most significantly required upon linezolid treatment. List of 25 genes most differentially required (*P* value <0.0005) under linezolid treatment as determined by TnSeq sorted by *P* values derived from permutation test and catalogued by predicted function. Adjusted *P* values are corrected for multiple tests using the Benjamini-Hochberg procedure.

### Membrane transporter knockdown increases susceptibility to linezolid

With the exception of *MAB_4116c*, which is involved in the export of cell envelope glycopeptidolipids^34^, the functions of these putative membrane transporters and their roles in drug resistance have yet to be characterized in *M. abscessus*. Thus, we first chose to examine the requirement for each of these transmembrane encoding genes upon linezolid treatment by utilizing an anhydrotetracycline-(ATc) inducible CRISPR interference (CRISPRi) system in the *M. abscessus* reference strain to generate transcriptional knockdowns^30,35^. Knockdown of each of these genes results in increased susceptibility to linezolid (Fig. 2a-i, Extended Data Fig. 3a-j), suggesting that each gene plays some role in determining linezolid efficacy. For the two *mmpL* genes identified in our genetic screen, *MAB_2303* and *MAB_4116c*, knockdown of their cognate *mmpS* membrane genes, *MAB_2302* and *MAB_4117c*, also led to increased linezolid sensitivity (Extended Data Fig. 3i,j). These genes did not appear as significant hits in the original screen, most likely due to their small size, which results in too few potential transposon insertion sites. Together these data suggest that each of these membrane transporters plays a role in establishing intrinsic linezolid resistance.

**Figure 2:**
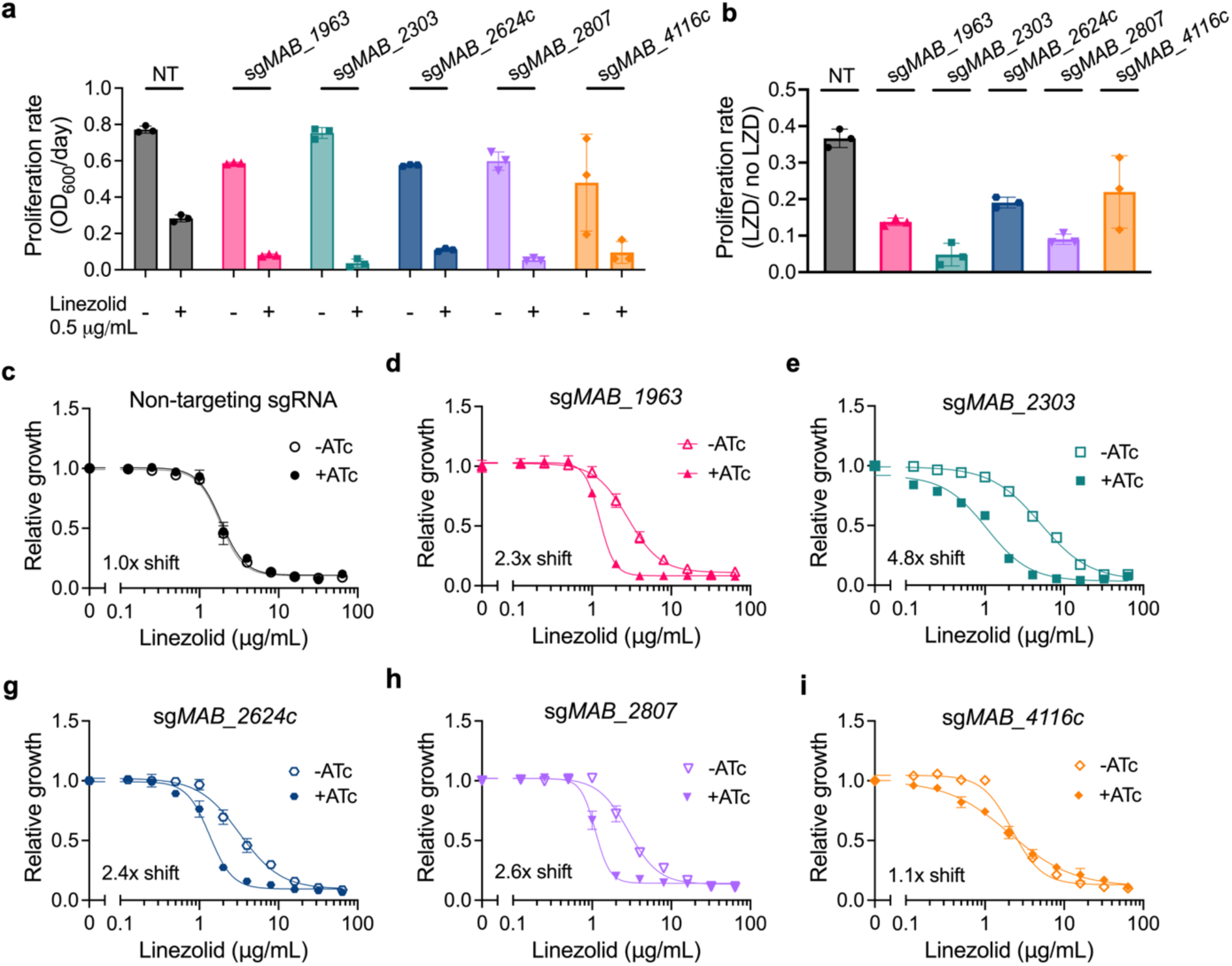
Knockdown of membrane transporters increases sensitivity to linezolid. **a,** Proliferation rates of *M. abscessus* ATCC19977 CRISPRi strains pre-depleted by treatment with 500 ng mL^−1^ ATc for 24 hours and then treated with 0.5 μg mL^−1^ linezolid or vehicle for 48 hours. Proliferation rates calculated from optical density of cultures over time. **b,** Ratio of proliferation rates of pre-depleted *M. abscessus* ATCC19977 CRISPRi strains treated with 0.5 μg mL^−1^ linezolid or vehicle along with ATc for 48 hours. Data are represented as individual values along with mean ± s.d. n=3 biological replicates. **c-i,** Relative growth of pre-depleted *M. abscessus* ATCC19977 strains as measured by reduction of a colorimetric dye after treatment with indicated concentrations of linezolid along with ±500 ng mL^−1^ ATc for 24 hours. Values normalized to vehicle only control per strain. All data are represented as individual values along with mean ± s.d. n=3 biological replicates. NT = non-targeting sgRNA. KD = knockdown. ATc = anhydrotetracycline. LZD = linezolid.

### Candidate genes impact both general cell permeability and efflux

We next interrogated the mechanistic role of each of these proteins in mediating linezolid susceptibility. Several of the knockdown strains display baseline growth defects (Fig. 2a-b, Extended Data Fig. 3a-h), suggesting that they play an important role in the normal biology of the bacterium. However, knockdown of these genes does not induce any gross alterations to general cell morphology (Extended Data Fig. 4), indicating that these genes’ effects on linezolid susceptibility are not mediated by substantial cellular deformities. Instead, we posited that these membrane proteins may have specific effects on drug uptake. We first tested if these knockdown strains have altered general permeability to chemicals by measuring accumulation of calcein AM, a hydrophobic, non-fluorescent molecule that can passively diffuse through cells; once inside cells, calcein AM is cleaved by host esterases into calcein, a fluorescent product, that remains intracellular^36^. Knockdown of *MAB_2303*, *MAB_2624c*, or *MAB_2807* does not induce increased intracellular calcein accumulation, suggesting that knockdown of these genes does not alter general permeability of the cell (Fig. 3a, Extended Data Fig. 5a-h). However, *MAB_1963* and *MAB_4116c* knockdowns display increased intracellular calcein accumulation, suggesting that these genes play a role in limiting the general permeability of the cell (Fig. 3a, Extended Data Fig. 5a-h). Furthermore, these results are consistent with previous literature that links *MAB_4116c* with production of glycopeptidolipids, which have been implicated in establishment of the barrier function of *M. abscessus*^34,37^. However, what role *MAB_1963* may play in general cell permeability is currently unknown.

**Figure 3:**
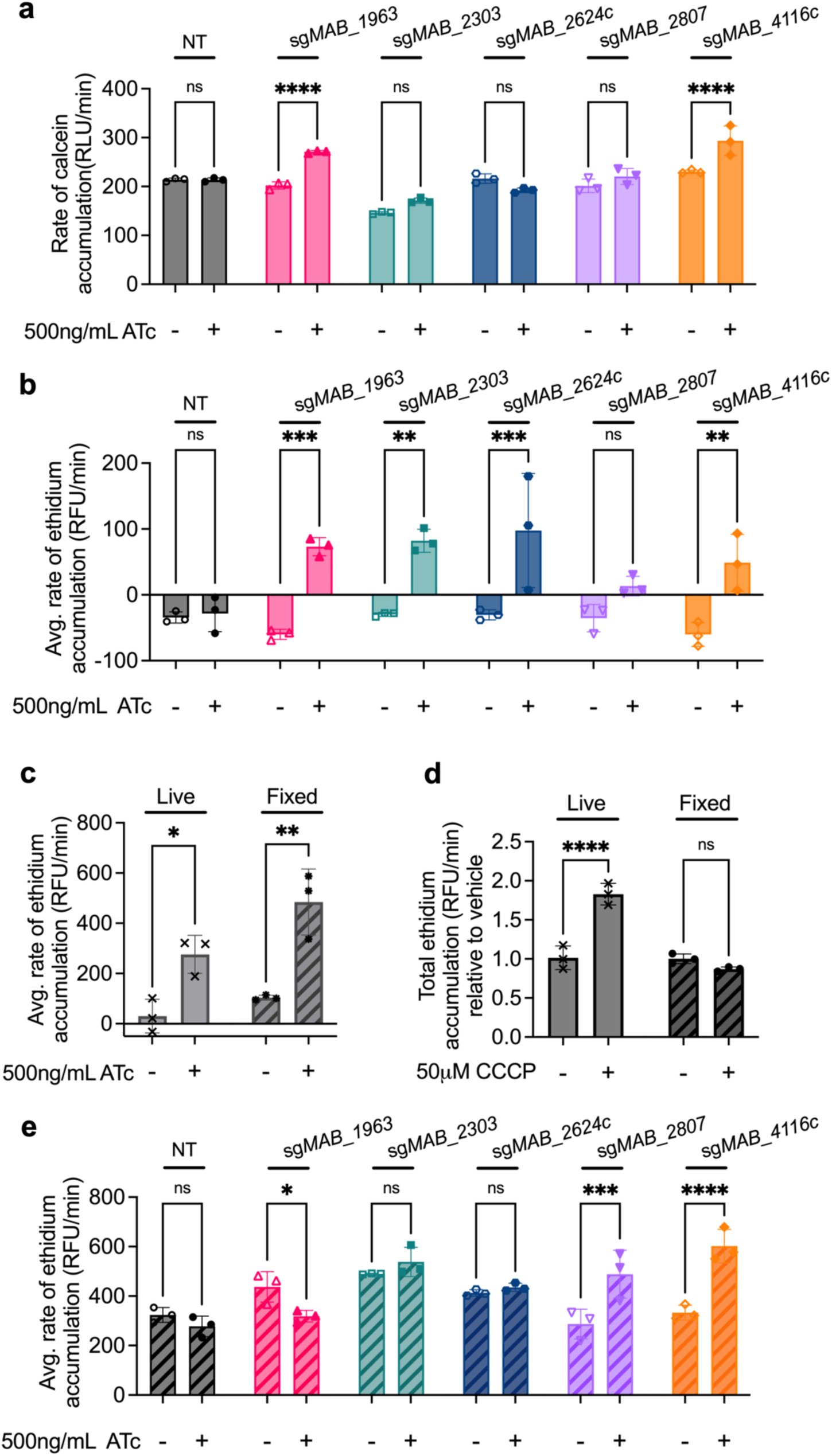
Membrane transporters disrupt general cell permeability or efflux. **a,** Rate of calcein accumulation as measured by calcein fluorescence in CRISPRi strains pre-treated with 500ng mL^−1^ ATc for 24 hours prior to addition of calcein AM. Values normalized to vehicle only control per strain. **b,** Rate of ethidium accumulation as measured by fluorescence in live membrane transporter CRISPRi strains pre-treated with 500ng mL^−1^ ATc for 24 hours prior to addition of ethidium bromide in. **c,** Rate of ethidium accumulation as measured by fluorescence in strains pre-treated with 500ng mL^−1^ ATc for 24 hours prior to addition of ethidium bromide in live and fixed PBP-lipo CRISPRi strain. **d**, Rate of ethidium accumulation as measured by fluorescence in live and fixed *M. abscessus* ATC19977 wildtype treated with 50μM CCCP. **e,** Rate of ethidium accumulation as measured by fluorescence in live and fixed membrane transporter CRISPRi strains pre-treated with 500ng mL^−1^ ATc for 24 hours prior to addition of ethidium bromide. All values normalized to vehicle only control per strain. Data for all graphs are represented as individual values along with mean ± s.d. n=3 biological replicates. Statistical significance was calculated with two-way ANOVA; \**P* < 0.05, \*\**P* < 0.01, \*\*\**P* < 0.001, \*\*\*\**P* < 0.0001. ns = not significant. NT = non-targeting sgRNA. ATc = anhydrotetracycline. CCCP = carbonyl-cyanide m-chlorophenylhydrazone.

To further examine general permeability as well as efflux, we performed an ethidium accumulation assay. Ethidium is an efflux substrate that increases in fluorescence when intercalated with DNA^38^. Either high cell permeability or low efflux rate will result in ethidium accumulation within the cell, leading to increased fluorescence intensity over time^38^. Knockdown of *MAB_1963*, *MAB_2303*, *MAB_2624c*, and *MAB_4116c* all lead to significantly increased rates of ethidium accumulation (Fig. 3b, Extended Data Fig. 5i-p), suggesting that these genes play a role either in establishing cell permeability or efflux of ethidium.

To distinguish between these possibilities, we compared ethidium accumulation in live and fixed cells. We reasoned that fixation would preserve the gross architecture of the cell wall but cease metabolism and thus eliminate the influence of active efflux on ethidium accumulation. Indeed, upon knockdown of PBP-lipo, a cell wall synthesis gene previously shown to increase membrane permeability to hydrophobic compounds^30^, both live and fixed cells exhibited significantly increased rates of ethidium accumulation (Fig. 3c, Extended Data Fig. 6s,t), demonstrating that disruption to membrane permeability is preserved in fixed cells. In contrast, efflux pumps that do not disrupt general membrane permeability should only impact ethidium accumulation in live cells, but not fixed. Consistent with this model, total ethidium accumulation was significantly increased in live cells treated with carbonyl-cyanide m-chlorophenylhydrazone (CCCP), a protonophore that disrupts the proton gradient and thus hinders efflux activity^39^, but was unchanged in fixed cells treated with CCCP (Fig. 3d, Extended Data Fig. 5q,r). Thus, genes that impact ethidium uptake in live, but not fixed cells are likely involved in efflux.

Using this methodology, we examined whether each membrane protein of interest is involved in efflux, general permeability, or both. Knockdown of *MAB_4116c* and *MAB_2807* still resulted in increased rates of ethidium accumulation after fixation (Fig. 3e, Extended Data Fig. 5u-ab), suggesting that general cell wall permeability is likely compromised in those strains, allowing for greater ethidium accumulation despite the cessation in cell metabolism. In contrast, knockdown of *MAB_2303* and *MAB_2624c* did not lead to increased rates of calcein accumulation (Fig. 3e) and only altered rates of ethidium accumulation in live cells (Fig. 3c,e), suggesting that these genes may play a role in active efflux.

### MAB_2303 is an efflux pump that plays a direct role in linezolid efflux

Given the substantial increase in linezolid susceptibility induced by *MAB_2303* knockdown (Fig. 2), we chose to characterize this gene further. Based on the calcein and ethidium accumulation assays (Fig. 3), we posited that MAB_2303 may directly efflux linezolid, leading to drug resistance. To rule out the possibility that knockdown of *MAB_2303* disrupts membrane or cell wall biology through indirect alteration of protein complex formation, we generated an enzymatically dead mutant of MAB_2303 through a tyrosine-444 to histidine (Y444H) mutation predicted to abrogate function^40–42^. The resulting protein should participate in protein complexes but fail to perform its enzymatic function. Expression of an sgRNA-resistant construct containing both *MAB_2302* and *MAB_2303* to ensure proper expression of both members of the operon was able to rescue linezolid sensitivity of the *MAB_2303* knockdown (Fig. 4a-b). In contrast, the Y444H mutant was only partially able to rescue linezolid sensitivity (Fig. 4a-b), indicating that enzymatic function of MAB_2303 is required to fully counteract the effects of linezolid.

**Figure 4.**
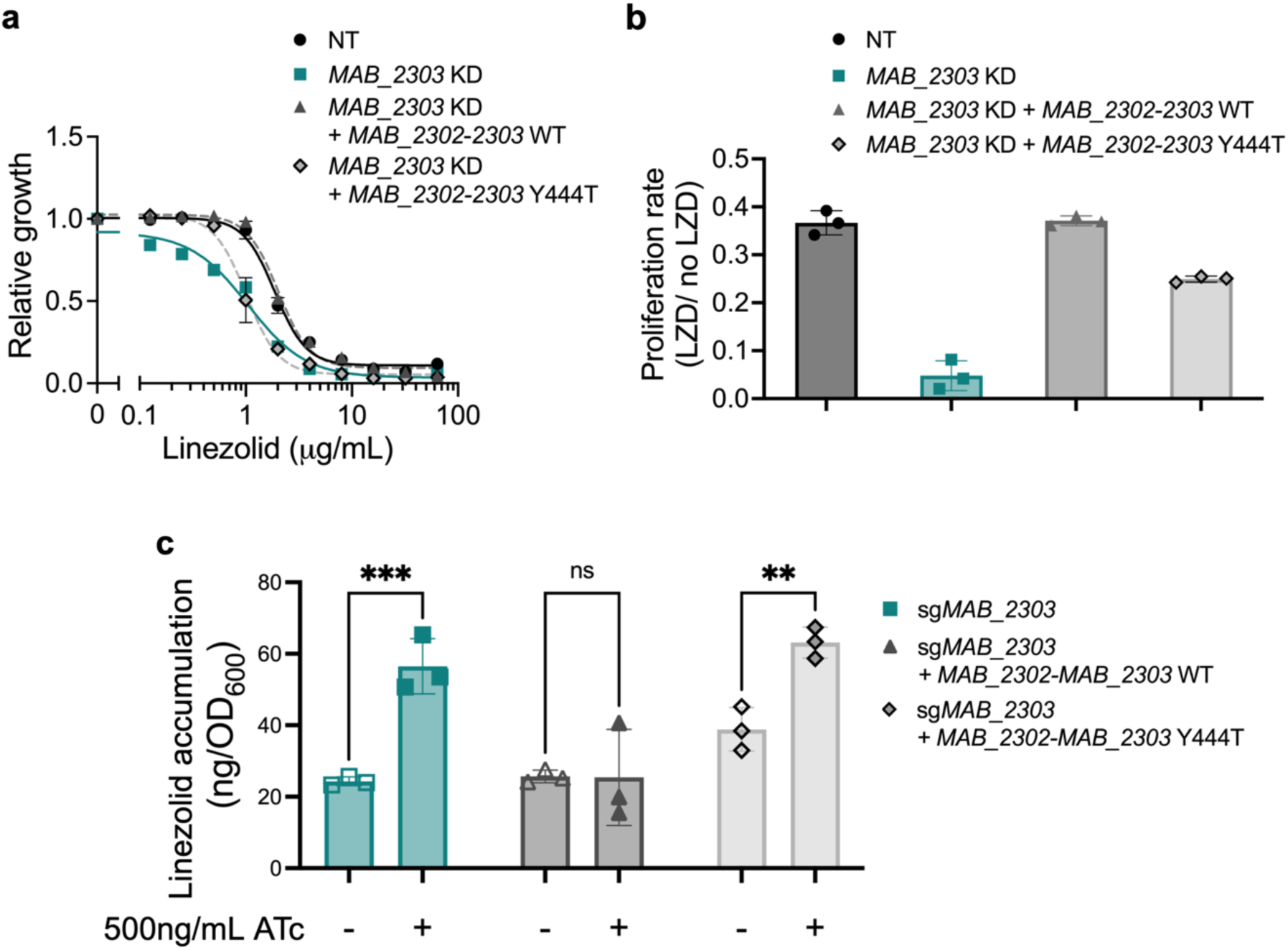
MAB_2303 effluxes linezolid. **a,** Relative growth of *M. abscessus* with non-targeting sgRNA (NT) or sg*MAB_2303* complemented with either wildtype (WT) or mutant (Y444T) sgRNA-resistant *MAB_2302-2303* treated with indicated concentrations of linezolid in the presence or absence of 500 ng mL^−1^ ATc for 48 hours. Values normalized to vehicle only control for each strain. **b,** Ratio of proliferation rates of pre-depleted *MAB_2303* knockdown strains complemented with either wild type (WT) or mutant (Y444T) sgRNA-resistant *MAB_2302-2303* treated with 0.5 μg mL^−1^ linezolid in the presence or absence of 500ng mL^−1^ ATc for 48 hours. Proliferation rates calculated from optical density of cultures over time. **c,** LC-MS measurement of intracellular linezolid accumulation in pre-depleted *MAB_2303* knockdown strains complemented with either wild type (WT) or mutant (Y444T) sgRNA-resistant *MAB_2302-2303*. Values are normalized to initial OD_600_ measurements. All data are represented as individual values along with mean ± s.d. n=3 biological replicates. Statistical significance was calculated with two-way ANOVA; \**P* < 0.05, \*\**P* < 0.01, \*\*\**P* < 0.001, \*\*\*\**P* < 0.0001. ATc = anhydrotetracycline. NT = non-targeting sgRNA. KD = knockdown. LZD = linezolid.

To examine directly whether linezolid accumulation is affected by MAB_2303, we measured the intracellular accumulation of linezolid in the induced knockdown mutants by liquid chromatography-mass spectrometry (LC-MS). *MAB_2303* knockdown increases linezolid accumulation in cells, and that accumulation is rescued by constitutive expression of the *MAB_2302-2303* operon (Fig. 4c, Supplemental Table 5), supporting the role of MAB_2303 as a linezolid efflux pump. Further, expression of the *MAB_2302-2303* Y444H operon fails to reduce linezolid accumulation (Fig. 4c, Supplemental Table 5), consistent with its inability to rescue linezolid susceptibility and supporting the conclusion that MAB_2303 plays a direct and active role in the efflux of linezolid.

### *MAB_2303* only alters susceptibility to compounds chemically similar to linezolid

If MAB_2303 is responsible for exporting linezolid, its knockdown might alter susceptibility to other antibiotics with similar chemical structures. To identify compounds that might be exported by MAB_2303, we clustered 529 antibiotics based on their chemical similarity both by atom pair similarity (Fig. 5a) and a fragment-based approach (Fig. 5b) and identified the clinically relevant antibiotics pretomanid, chloramphenicol, and bedaquiline as chemically similar compounds that might be subject to efflux by MAB_2303. Indeed, accumulation measurements of 19 structurally diverse antibiotics in *M. abscessus* indicated that MAB_2303 limits accumulation of pretomanid, chloramphenicol, and trimethoprim in addition to linezolid (Fig. 5c). In contrast, MAB_2303 did not prevent uptake of compounds structurally divergent from linezolid, such as clarithromycin and rifampicin. Together, these results suggest that antibiotic efflux follows predictable chemical patterns, and that MAB_2303 may provide resistance to antibiotics with similar structure to linezolid.

**Figure 5:**
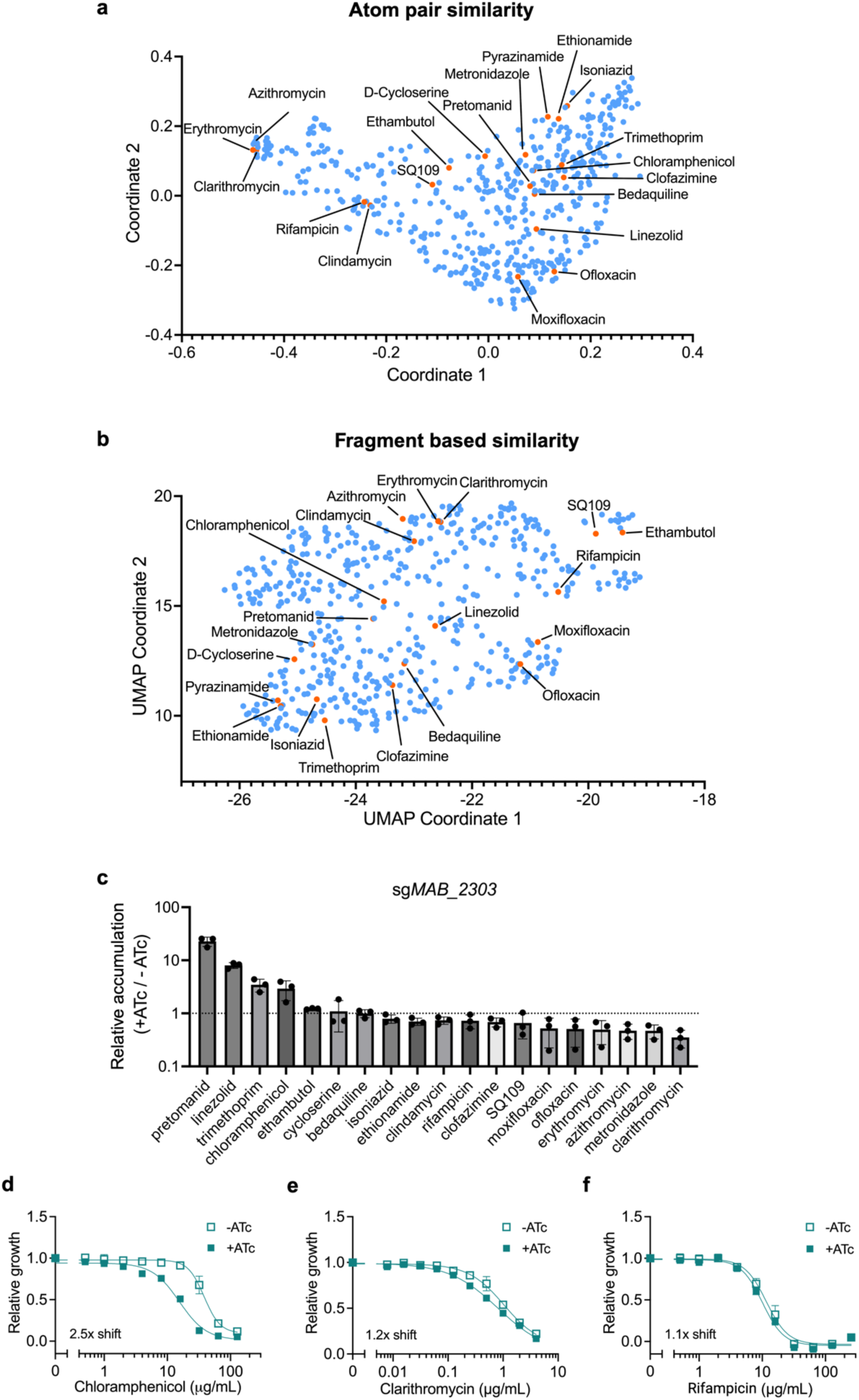
MAB_2303 effluxes compounds that share similar chemical properties. **a-b,** 529 antibiotics represented by **(a)** multidimensional scaling of atom pair similarity and **(b)** UMAP dimensional reduction of fragment-based similarity. Panel of mycobacterial antibiotics from Fig. 1 are highlighted in orange and labeled. **c,** LC-MS measurement of intracellular accumulation of indicated antibiotics in pre-depleted (+ATc) sg*MAB_2303* CRISPRi *M. abscessus* strain incubated for 4 hr with antibiotics. Values normalized to internal standard, initial antibiotic levels in media prior to incubation, and to a non-depleted strain control (-ATc). Data are represented as individual values along with mean ± s.d. n=3 biological replicates. **d-f,** Relative growth of pre-depleted *MAB_2303* knockdown strain as measured by reduction of a colorimetric dye after treatment with indicated concentrations of **(d)** chloramphenicol, **(e)** clarithromycin, and **(f)** rifampicin in the presence or absence of 500 ng mL^−1^ ATc for 24 hours. Values normalized to vehicle only control per drug. Data are represented as individual values along with mean ± s.d. n=3 biological replicates. ATc = anhydrotetracycline.

To test this hypothesis, we examined whether MAB_2303 knockdown alters chloramphenicol efficacy, as *M. abscessus* is not responsive to pretomanid^43^ or trimethoprim^44^, precluding determination of susceptibility to those compounds. Consistent with the model, knockdown of *MAB_2303* increases susceptibility to chloramphenicol (Fig. 5d), but does not affect susceptibility to structurally dissimilar compounds such as clarithromycin and rifampicin (Fig. 5e,f, Extended Data Fig. 6a-e). Further, the lack of effect on susceptibility to ribosome-targeting antibiotics, including clarithromycin and amikacin (Extended Data Fig. 6a), suggests that MAB_2303 does not have direct effects on the ribosome. Of interest, MAB_2303 knockdown does result in a modest increase in susceptibility to fluoroquinolone antibiotics (Extended Data Fig. 6d,e) without increasing their accumulation, suggesting that MAB_2303 may play other roles in intrinsic antibiotic resistance beyond direct drug efflux. Together, these results indicate that MAB_2303 contributes to the *M. abscessus* permeability barrier within a relatively specific chemical space and argues that drug impermeability and lack of accumulation is likely driven by the overlapping effects of numerous such membrane proteins.

## Discussion

An impermeable cell wall and active efflux of drugs have long been thought to be among the most important components of mycobacteria’s arsenal against antibiotics^6^. In keeping with that model, this work demonstrates that antibiotic resistance positively correlates with antibiotic permeability and retention, suggesting that a combination of drug efflux and an impermeable cell wall likely plays a major role in intrinsic antibiotic resistance in *M. abscessus*. Several membrane transporters across superfamilies have been demonstrated in *M. tuberculosis* (Mtb) and *Mycobacterium smegmatis* to maintain low chemical permeability and efflux certain antibiotics,^13,14^ but most membrane transporters remain largely undefined in *M. abscessus*. Despite the lack of characterization of these proteins, *M. abscessus* does appear to possess machinery that could be responsible for promoting a high degree of cell envelope integrity and drug efflux. For example, *M. abscessus* possesses 31 putative MmpL proteins^45^ compared to Mtb with only 13^46^. Consistent with this possibility, we have now identified several *M. abscessus*-specific putative membrane proteins, including two MmpL proteins, that play a role in cell wall permeability or drug efflux against linezolid. Additionally, most of these genes lack significant shared homology with the closest relative Mtb orthologs, reinforcing the idea that efficient drug efflux and impermeability may be responsible for the elevated level of broad-spectrum drug resistance observed in *M. abscessus* compared to other mycobacteria.

Membrane transporters and efflux pumps are often thought to display a high degree of redundancy, which makes it challenging to identify and interrogate the role of these proteins in drug resistance. Our study suggests that while these proteins may have overlapping impacts on drug uptake, they appear to carry out these roles in mechanistically distinct ways. For example, this work demonstrates that several transmembrane proteins from three of the five superfamilies of transmembrane proteins all individually play a role in linezolid susceptibility. Given the diversity in these transmembrane proteins, their functions, as well as their native substrates, likely vary. This suggests that multiple pathways contribute to membrane integrity and independently reduce linezolid susceptibility. Future work into these proteins’ native substrates could illuminate how these transporters limit susceptibility to linezolid and other antibiotics. This work underscores the concept that targeting membrane proteins could provide attractive options for augmenting current antibiotic therapy regimens, but that those targets should be carefully designed to maximize the uptake of the desired antibiotics.

We have characterized a novel *M. abscessus*-specific MmpL protein, MAB_2303, as a linezolid drug efflux pump, and identified several additional MmpL and MmpS proteins that appear to play a role in linezolid resistance. Drug design for targeting essential MmpL proteins is an area of active research and several potential MmpL inhibitors in *M. smegmatis* and *M. tuberculosis* have been identified^47^, suggesting that these proteins might be inhibited with new drugs. While most research on MmpL inhibitors has focused on targeting essential MmpL proteins like MmpL3^48^, our work suggests that identifying compounds that broadly target non-essential MmpL proteins could be an attractive potentiating strategy for combating intrinsic antibiotic resistance.

Additionally, we find that antibiotic substrates exported by an efflux pump can be computationally predicted through several methods of chemical similarity scoring and validated with a streamlined LC-MS approach. Through this approach we found that MAB_2303 likely is responsible for the efflux of other chemically similar compounds, including chloramphenicol, pretomanid and trimethoprim, but not for efflux of chemically dissimilar compounds. It remains unclear how MmpL proteins distinguish between different substrates^9,49^; however, our data, as well as that of others, suggest that antibiotics or endogenous substrates that share similar chemical properties, including size, charge, and hydrophobicity tend to be substrates of the same MmpL protein^9,49^. Further research in how MmpL proteins choose substrates can help us to better understand how they also allow for expulsion of certain antibiotics. Additionally, given the interest in targeting MmpL proteins for potentiating antibiotics, a more thorough understanding of their biology could aid in rational drug design that could disrupt efflux of multiple antibiotics with shared chemical properties.

Together, these results suggest that low antibiotic uptake and retention are important intrinsic antibiotic resistance mechanisms in *M. abscessus*. Future work to target membrane proteins involved in intrinsic antibiotic resistance could provide therapeutic avenues to increase efficacy of current antibiotics and improve the outcomes for *M. abscessus* infections.

## Acknowledgements

We thank the Harvard Center for Mass Spectrometry for assistance with LC-MS experiments and the Biopolymers Facility at Harvard Medical School for sequencing. T.F. was supported by a Boehringer Ingelheim Fonds MD fellowship. M.R.S. received support as a Merck Fellow of the Damon Runyon Cancer Research Foundation, DRG-2415-20. E.J.R. was supported by NIH/NIAID under award number R01AI179642. Figures created with BioRender.com.

## Materials Availability

All reagents generated in this study are available upon request from the corresponding author.

## Data Availability

All relevant data generated in this study are present within the manuscript and Supplemental Information except for raw TnSeq data, which will be available on SRA. Project and accession numbers will be listed prior to publication.

## Materials and methods

### Strains

All experiments were performed in the *Mycobacterium abscessus subspecies abscessus* type strain (ATCC19977) unless otherwise indicated. Clinical isolate *M. abscessus subspecies massiliense* BWH-F was isolated from a skin biopsy^31^. All plasmid construction was performed in DH5α *Escherichia coli*.

### Mycobacterial culturing conditions

*M. abscessus* liquid cultures were grown in Middlebrook 7H9 broth (271310, BD Diagnostics) with 0.2% (v/v) glycerol (GX0185, Supelco), 0.05% (v/v) Tween-80 (P1754, MilliporeSigma), and 10% (v/v) oleic acid-albumin-dextrose-catalase (OADC) (90000-614, VWR) or 10% (v/v) albumin-dextrose-catalase (ADC) composed of 50 g/L albumin, 0.03 g/L catalase, 8.5 g/L NaCl, and 20 g/L dextrose. Cultures were shaken at 150 r.p.m. at 37°C.

### Mycobacterial transformations

*M. abscessus* strains were grown to an optical density (OD_600_) of 0.8, and washed thrice with sterile 10% glycerol by pelleting at 5000 x *g* for 7 minutes at 22°C. After final wash, cells were resuspended in 1% of the initial culture volume in 10% glycerol. 50 µL of electrocompetent mycobacteria were mixed well with 100 ng plasmid in 2 µL water and then transferred to a 2 mm electroporation cuvette (89047-208, VWR). The cells were electroporated at 2500 V, 125 Ω, 25 μF using an ECM 630 electroporator (45-0651, BTX). 1 mL 7H9 +OADC broth was added to the electroporated cells, and cells recovered for 4 hours shaking at 150 r.p.m. at 37°C. 100 µL of recovered cells were spread on 7H10 + 0.5% (v/v) glycerol + 10% (v/v) OADC agar plates with 50 µg mL^−1^ kanamycin sulfate using 4 mm borosilicate glass beads. Plates were incubated at 37°C for 4 days or until colonies were visible.

### Generation of CRISPRi and CRISPRi-resistant mutant strains

CRISPRi plasmids were constructed as previous described^35^ using Addgene plasmid 166886. Briefly, plasmid 166886 was digested overnight at 55°C with BsmBI-v2 (R0739L, New England BioLabs) and then gel purified (T1020, New England BioLabs). Three sgRNAs were selected using Pebble sgRNA Design Tool (Rock Lab, Rockefeller University) to target three different locations of the non-template strand of each gene of interest. Each individual sgRNA with appropriate overhangs was annealed and ligated using T4 ligase (M0202M, New England BioLabs) into three separate BsmBI-v2 digested backbones. To generate a triple CRISPRi plasmid with all three sgRNAs, SapI-based Golden Gate cloning site 3′ to the first sgRNA scaffold was used as previously described^35^. CRISPRi NT was constructed in a similar manner but with scrambled, non-targeting sgRNAs. Successful plasmid construction was verified using whole plasmid sequencing with Oxford Nanopore Technologies (ONT) (Plasmidsaurus). Triple CRISPRi plasmids were transformed into ATCC19977 as described above and selected on 7H10 + 0.5% (v/v) glycerol + 10% (v/v) OADC agar plates containing 50 µg mL^−1^ kanamycin sulfate.

*MAB_2302-MAB_2303* rescue plasmids were constructed by introducing synonymous mutations at the protospacer adjacent motif (PAM) and seed sequences for all sites of sgRNA targeting. Additionally, a second version of the *MAB_2302-MAB_2303* rescue plasmid was constructed with the same CRISPRi-resistant synonymous mutations as well as a nonsynonymous mutation (Y444H) at the proposed catalytic site in the *mmpL* (*MAB_2303*). Gene fragments (Azenta) containing these synonymous mutations and/or a nonsynonymous mutation in the catalytic site with NdeI (R0111S, New England Biolabs) and XhoI (R0146S, New England Biolabs) overhang sites were restriction enzyme digested and then assembled into Tweety-integrating zeocin marked MOP plasmids using the Gibson Assembly standard protocol (E5510, New England Biolabs). Successful plasmid construction was verified using whole plasmid sequencing with Oxford Nanopore Technologies (ONT) (Plasmidsaurus). Plasmids transformed into the CRISPRi sg*MAB_2303* strain using transformation protocol described above and selected on 7H10 + 0.5% (v/v) glycerol + 10% (v/v) OADC agar plates containing 50 µg mL^−1^ zeocin and 50 µg mL^−1^ kanamycin sulfate.

### Minimum inhibitory concentration determination

*M. abscessus* strains were grown until mid-log phase (OD_600_ of 0.6-0.8). Strains were induced for knockdown 18-24 hours prior to start of the assay with 500 ng μL^−1^ ATc. Cultures were then diluted to OD_600_ of 0.003 and 200 μl aliquots were plated in biological triplicate in wells (3370, Corning) containing specified antibiotics as well as fresh 500 ng μL^−1^ ATc or vehicle when relevant. Antibiotic stocks were made as follows: 20 mg mL^−1^ linezolid (PZ0014, Sigma-Aldrich) in DMSO, 10 mg mL^−1^ clarithromycin (C9742, Sigma-Aldrich) in DMSO, 10 mg mL^−1^ amikacin disulfate salt (A1774, Sigma-Aldrich) in water, 20 mg mL^−1^ rifampicin (R3501, Sigma-Aldrich) in DMSO, and 10 mg mL^−1^ moxifloxacin (SML1581, Sigma Aldrich) in DMSO. The cells were then incubated at 37°C with shaking at 150 r.p.m. for 24 hours. 0.002% resazurin (R7017, Sigma Aldrich) in ddH2O was spiked into each well and plates were incubated for an additional 24 hours at 37°C with shaking at 150 r.p.m. MIC determination was conducted using a Tecan Spark 10M plate reader (Mannedorf, Switzerland) by measuring absorbance at 570nm and 600nm and normalizing the ratio to background and no drug control.

Minimum inhibitory concentrations for Fig. 1c were derived from prior literature: clofazimine^50^, azithromycin^51^, bedaquiline^52^, clarithromycin (this study), erythromycin (this study), linezolid (this study), moxifloxacin (this study), chloramphenicol^53^, SQ109^54^, ofloxacin (this study), ethambutol^2^, pretomanid^55^, metronidazole^55^, cycloserine^56^, clindamycin^51^, rifampicin (this study), isoniazid^57^, and trimethoprim^44^.

### Growth curve

CRISPRi strains were grown until mid-log phase (OD_600_ of 0.6-0.8) and then pre-depleted with ATc at 500 ng mL^−1^ for 18-24 hours. Cultures were then back-diluted at final OD_600_ of 0.02 and 200 µl of diluted cells were added in biological triplicates with DMSO or 0.5 µg mL^−1^ linezolid and fresh ATc at 500 ng mL^−1^ when relevant. Growth was determined by continuous OD_600_ measurement in 15-minute intervals in a Spark 10M plate reader for 48 hours at 37°C with continuous shaking at 1000 rpm. Growth curve data were analyzed using Microsoft Excel 365 and GraphPad Prism 9.

### Transposon library production

The BWH-F transposon mutant library (69.5% TA insertion coverage) was previously generated^31^. Briefly, the *M. abscessus* BWH-F strain was transduced with temperature-sensitive φMycoMarT7 phage carrying the Himar1 transposon. After selection with 100µg mL^−1^ kanamycin, mutant libraries with >150,000 individual bacterial mutants were harvested and stored in aliquots with 7H9 + 10% (v/v) glycerol at −80°C.

### Transposon library growth conditions and selection

Transposon mutant BWH-F library was inoculated in biological triplicates at 2.1×10^7^ CFU mL^−1^ into 7H9 + 0.5% (v/v) glycerol + 10% (v/v) OADC either incubated with 16 µg mL^−1^ linezolid or DMSO. After 11 doublings, cultures were pelleted at 5,000 x *g* for 5 min at 22°C, resuspended in 7H9 + 0.2% (v/v) glycerol + 0.05% (v/v) Tween-80 + 10% (v/v) OADC, mixed equal volume with 50% glycerol, and frozen at −80°C. The harvested replicates were then titered by plating on 7H10 + 0.5% (v/v) glycerol + 10% (v/v) OADC agar plates supplemented with 100 µg mL^−1^ kanamycin sulfate. 150,000 cells of each replicate were plated 7H10 + 0.5% (v/v) glycerol + 10% (v/v) OADC + 0.005% Tween 80 + 100 µg mL^−1^ kanamycin sulfate on six 245 mm x 245 mm plates (431111, Corning). Colonies were grown for 4 days at 37°C. Each replicate was combined via scraping into a 50 mL conical tube containing 5 ml 7H9 + 0.2% (v/v) glycerol + 0.05% (v/v) Tween-80 + 10% (v/v) OADC and 5 mL 50% glycerol. 2 mL aliquots of the libraries were stored at - 80°C for genomic DNA extraction.

### Genomic DNA Isolation

gDNA was isolated using an established protocol with minor adaptations^28,59^. The post-selection mutant libraries were thawed, pelleted at 5,000 x *g* for 5 min at 22°C and then resuspended in TE Buffer (10mM Tris HCl pH 7.4 and 1mM EDTA pH 8). Cell suspensions were transferred to 2 mL tubes containing 0.1 mm silica beads (116911500, MP Biomedicals) and 700 µL of 25:24:1 phenol:chloroform:isoamyl alcohol (P3803, MilliporeSigma). Bacteria were lysed utilizing a Bead Bug 3 Microtube Homogenizer (D1030, Benchmark Scientific, Sayreville, NJ, USA) four times at 45 second intervals at 4000 r.p.m. Samples were chilled on ice for 45 seconds in between cycles. Post-homogenization, cell debris was pelleted at 21,130 x *g* for 10 minutes at 22°C. The aqueous phase was combined with an equal volume of 25:24:1 phenol:chloroform:isoamyl alcohol and incubated on a rocker for 1 hour at 22°C. The mixtures were then transferred to pre-pelleted MaXtract High-Density phase-lock tubes (129065, Qiagen), followed by centrifugation at 1500 x *g* for 5 minutes at 4°C. ½ volume of chloroform (193814, MP Biomedicals) was added to upper aqueous phase and centrifuged at 1500 x *g* for 5 minutes at 4°C.The upper aqueous layers were transferred to new MaXtract High-Density phase-lock tubes and incubated with RNase A at 25 µg mL^−1^ (EN0531, Thermo Fisher Scientific) at 150 r.p.m. for 1 hour at 37°C. Samples were re-extracted with an equal volume of 25:24:1 phenol:chloroform:isoamyl alcohol, followed by centrifugation at 1500 x *g* for 5 minutes at 4°C. A second extraction with ½ volume chloroform was performed, and centrifuged at 1500 x *g* for 5 min at 4°C. The aqueous phase containing the DNA of each sample was transferred to a fresh conical tube prepared with 1/10th volume of 3M sodium acetate pH 5.2 (3032-16, VWR) and 1 volume of isopropanol (3032-16, VWR), followed by an overnight incubation at 22°C. The DNA pellets were washed thrice with 5 mL 70% ethanol at 5000 x *g* for 10 min, dried for 10 minutes to eliminate residual ethanol, and resuspended in 500 µL nuclease-free water. DNA concentration of the samples was quantified with Qubit Fluorometer (Q33238, Thermo Fisher Scientific) using the Broad Range assay kit (Q33266, Thermo Fisher Scientific).

### Transposon sequencing, mapping, and analysis

Sequencing libraries were generated from the isolated genomic DNA by amplifying chromosomal-transposon junctions, following an established protocol outlined by Long et al. 2015^60^ and sequenced on an Illumina NextSeq 500 sequencer. The resultant reads were aligned to the BWH-F genome (SRA project number PRJNA840944, accession SRX15416547). Analysis of the data was conducted employing TRANSIT^61^. Insertion counts at each TA site were subjected to trimmed total reads normalization and were then averaged across replicates. The resampling analysis in TRANSIT was applied for the comparative assessment of essentiality between genes in the no drug and linezolid exposed strains. TA site insertions at the 5% C- or N-terminus of each gene were trimmed. *P* values were derived from a permutation test, and adjusted *P* values for multiple tests were derived from the Benjamini-Hochberg method.

### Calcein accumulation assay

CRISPRi strains were grown until mid-log phase (OD_600_ of 0.6-0.8) and then back diluted to pre-deplete target proteins with ATc 500 ng mL^−1^ or DMSO for 18-24 hours. Bacteria were washed twice with 1x PBS pH 7.4 (10010049, Thermo Fisher) + 0.05% Tween80 (PBST). After final wash, cells were resuspended in 1x PBST for final OD_600_ of 0.4 and added to black-bottomed 96 well plate (07-200-722, Thermo Fisher). Plates were incubated at 37°C with shaking at 150 r.p.m. for 30 minutes. 1 µg mL^−1^ Calcein AM (C3100MP, Thermo Fisher) was then spiked into each well. Plates were incubated at 37°C with shaking in a Tecan Spark 10M plate reader with ex/em 488nm/520nm fluorescence every minute for 40 minutes.

### Ethidium accumulation assay

CRISPRi strains were grown until mid-log phase (OD_600_ of 0.6-0.8) and then back diluted to pre-deplete target proteins with ATc 500 ng mL^−1^ or DMSO for 18-24 hours. Bacteria were back diluted to an OD_600_ of 0.2 and half of the total volume was transferred directly to black-bottom 96-well plates (07-200-722, Thermo Fisher) or fixed with 2% (v/v) paraformaldehyde (sc-281692, ChemCruz) for 1 hour. Following fixation, cells were centrifuged at 5000 x *g* for 5 minutes and resuspended in the original volume of 7H9 media and transferred to black-bottom 96-well plates. Ethidium bromide (E7637, Sigma-Aldrich) was then added to a final concentration of 1 µg mL^−1^. The plate was shaken manually and incubated at 22°C for 10 minutes. Ethidium fluorescence was then measured at 530nm/600 nm ex/em in 1 minute cycles for 1 hour at 37°C using a Tecan Spark 10M plate reader.

### Arrayed antibiotic extraction

Wild type or CRISPRi strains were grown until mid-log phase (OD_600_ of 0.6-0.8) and then diluted back and cultured for 18-24 hours to pre-deplete target proteins with either 500 ng mL^−1^ ATc, DMSO, or no treatment for CRISPRi + ATC, -ATC, and wild type, respectively. Cultures were pelleted at 3200 x *g* for 10 minutes at room temperature then washed in 1X volume blood bank saline (89370-096, VWR) and resuspended in 7H9 + 0.5% (v/v) glycerol + 10% (v/v) ADC at a final OD_600_ of 15 with 500 ng mL^−1^ ATc or equivalent volume of DMSO added to appropriate cultures. Cells were added in biological triplicate to 96-well plates with 7H9 + 0.5% (v/v) glycerol + 10% (v/v) ADC containing a final concentration of 20 µM antibiotic in each of 20 wells resulting in a final OD_600_ = 7.5. 50 µL of media was immediately collected from each well and pooled, then filter sterilized through a 0.22 µm PVDF filter (SE1M179M6, Millipore). Media was stored at −80°C until extraction. Cells were incubated with antibiotics for 4 hr at 37°C with shaking at 150 rpm. After incubation, cultures were combined in a conical tube, then pelleted at 3200 x *g* for 10 minutes at 4°C and washed twice with pre-chilled blood bank saline. Pellets were resuspended in 0.8 mL 3:1:0.004 acetonitrile:methanol:formic acid + 10 nM verapamil, then transferred to 2 mL tubes containing 0.1 mm silica beads (116911500, MP Biomedicals). Bacteria were lysed utilizing a Bead Bug 3 Microtube Homogenizer (D1030, Benchmark Scientific, Sayreville, NJ, USA) three times at 45 second intervals at 4000 rpm. Samples were chilled on ice for 2 minutes in between cycles. Samples were pelleted at 21,130 x g for 10 minutes at 4°C. 50 µL of media was extracted with 450 µL 3:1:0.004 (v/v/v) acetonitrile:methanol:formic acid + 10 nM verapamil (V105, Millipore Sigma) as an internal standard. Media samples were vortexed for 1 minute at 22°C, then pelleted 10 minutes at 17,000 x g at 4°C. Supernatant from both media and cell pellet extractions was dried using a speedvac concentrator (Eppendorf 5305) 1 hr at 45°C, resuspended in 40 µL 3:1:0.004 (v/v/v) acetonitrile:methanol:formic acid, vortexed 1 minute at 22°C, and pelleted at 17,000 x g at 4°C. 35 µL supernatant was transferred to 9 mm plastic vials (Thermo C4000-11) with screw caps (Thermo 03-376-481) and stored at −80°C until LC-MS analysis.

### Linezolid extraction

CRISPRi strains were grown until mid-log phase (OD_600_ of 0.6-0.8) and then back diluted to pre-deplete target proteins with ATc 500 ng mL^−1^ or DMSO for 18-24 hours in biological triplicates in 50mL total volume each. Each replicate was normalized to an OD_600_ of 0.8 and the pelleted at 4000 x *g* for 10 minutes at room temperature then resuspended in 1/5^th^ the total volume with fresh 7H9 + 0.5% (v/v) glycerol + 10% (v/v) OADC. Fresh 500 ng mL^−1^ ATc or DMSO was added to appropriate cultures and then 20 µg mL^−1^ linezolid was added to each culture. Cultures were incubated for 2 hours at 37°C at 150 r.p.m. After incubation, cultures were pelleted at 4000 x *g* for 10 minutes at 4°C and washed once with pre-chilled 1x PBS. Pellets were stored at −80°C until metabolite extraction. Pellets were resuspended in pre-chilled 1mL of 2:2:1 acetonitrile (34851, Millipore Sigma) + methanol (439193, Millipore Sigma) + water with 2 µg mL^−1^ linezolid-d3 (25038, Cayman Chemicals) and 2 µg mL^−1^ verapamil (V105, Millipore Sigma). Cell suspensions were transferred 2 mL tubes containing 0.1 mm silica beads (116911500, MP Biomedicals). Bacteria were lysed utilizing a Bead Bug 3 Microtube Homogenizer (D1030, Benchmark Scientific, Sayreville, NJ, USA) for four times at 45 second intervals at 4000 r.p.m. Samples were chilled on ice for 2 minutes in between cycles. Post-homogenization, cell debris was pelleted at 21,130 x *g* for 10 minutes at 22°C.

### Liquid Chromatography-Mass Spectrometry

The LC-MS analysis for linezolid was following a reported method as described^58,59^. We applied an Agilent 1260 HPLC coupled with an Agilent 6120 quadrupole mass spectrum for compound accumulation analysis. Metabolites were separated using a 10-µL injection volume on a Chromolith SpeedRod RP-18 column (Sigma Aldrich) with a gradient of H_2_O (solvent A) and acetonitrile (solvent B) acidified with 0.1% formic acid. The gradient was as follows: 0 min, 10% B; 2 min, 10% B; 10 min, 100% B; 12 min, 10% B. Data were analyzed using Agilent ChemStation software to measure levels of the linezolid and d3-linezolid [M+H]^+^ ion with an accuracy of ± 20 ppm.,Signal intensity was quantified by standard curve of ratio of authentic linezolid versus d3-linezolid, and normalized by OD value at time of bacteria harvest.

LC-MS analysis for the antibiotic panel was performed using a QExactive+ orbitrap mass spectrometer (ThermoFisher) with a heated electrospray ionization (HESI) probe, coupled to a Dionex Ultimate 3000 UPLC system. 4 µL of extracted sample was injected into a Kinetex 2.6 µm EVO C18 column (150 x 2.1 mm), with the autosampler and column held at 4°C and 30°C, respectively. The chromatographic gradient consisted of 0.1% formic acid (solvent A) and 0.1% formic acid in acetonitrile (solvent B). The gradient was run as follows: 0-5 min: 1% solvent B, flow rate 0.3 mL/min; 5-15 min: linear gradient from 1-99% solvent B, flow rate 0.3 mL/min; 15-20 min: 99% solvent B, flow rate 0.3 mL/min; 20-25 min: 99% solvent B, flow rate 0.4 mL/min; 25-30 min: 1% solvent B, flow rate 0.3 mL/min. The mass spectrometer was operated in full scan, positive mode. The MS data acquisition was performed in a range of 100-1500 m/z, with the resolution set to 70,000, the AGC target at 1e6, and the maximum injection time at 50 msec.

After LC-MS analysis, antibiotic identification was performed with XCalibur 3.0.63 software (Thermo Fisher Scientific) using a 5ppm mass accuracy and a 0.5 min retention time window. Standards were used for assignment of antibiotic peaks at given m/z for the [M+H]^+^ ion and retention time and were compared to extraction buffer, media alone, and cells alone blanks. For metronidazole measurements, intracellular measurements represent hydroxymetronidazole. Linear range of detection was determined by analyzing four 10-fold dilutions of antibiotic standards containing 2 nmol, 200 pmol, 20 pmol, and 2 pmol of each antibiotic. Linear range was examined by ensuring that all data points fall within the 95% confidence interval of the linear regression, and by a Wald-Wolfowitz runs test to check deviation from linearity. Peak areas were normalized to verapamil internal standard and culture density. Relative antibiotic accumulation was calculated by normalizing intracellular antibiotic peak area after 4 hours of incubation to antibiotic peak area in media prior to incubation with *M. abscessus*.

### Antibiotic chemical similarity

529 antibiotics were selected for chemical similarity analysis by manually curating components of a commercially available antibiotic library (HY-L067 MedChemExpress, Princeton, NJ, USA). Atom pair similarity was calculated and represented using multidimensional scaling with ChemMineR v3.54.0^60^. Chemical fingerprints were calculated using FragFP in DataWarrior v06.02.01^61^ and represented as a 2-dimensional UMAP with nearest neighbors set to 100, minimum distance set to 0.5, and Euclidian distance.

---

**Extended Data Fig. 1:**
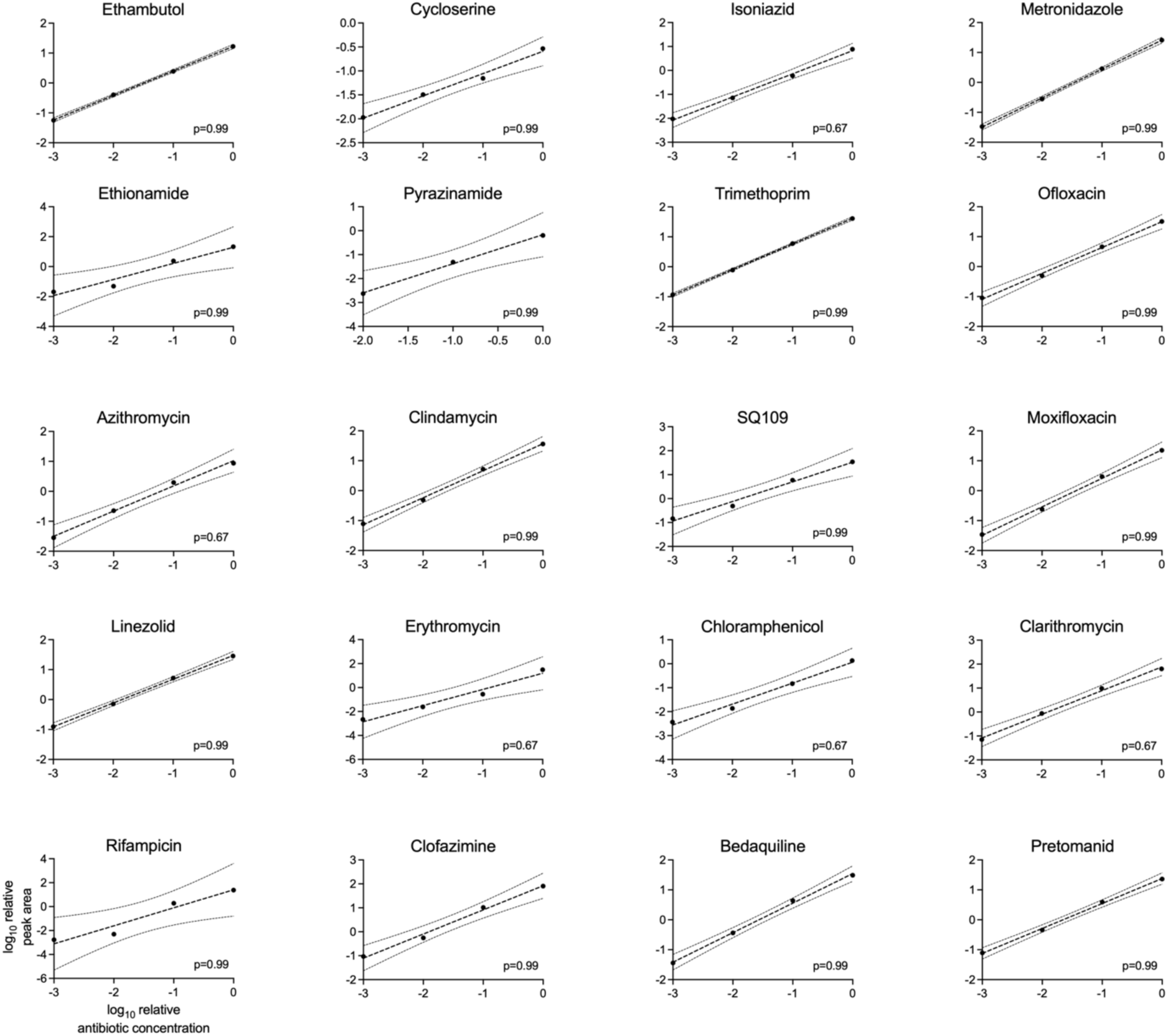
Antibiotic levels are measurable over a linear range. LC-MS measurement of indicated antibiotics over 1000-fold range of concentrations. Peak areas are normalized to internal standard, and antibiotic concentrations are normalized to the highest standard concentration. Line of best fit represents a simple linear regression and is represented +/− 95% confidence intervals. p-value derived from a Wald–Wolfowitz runs test to identify deviation from the linear fit.

**Extended Data Fig. 2:**
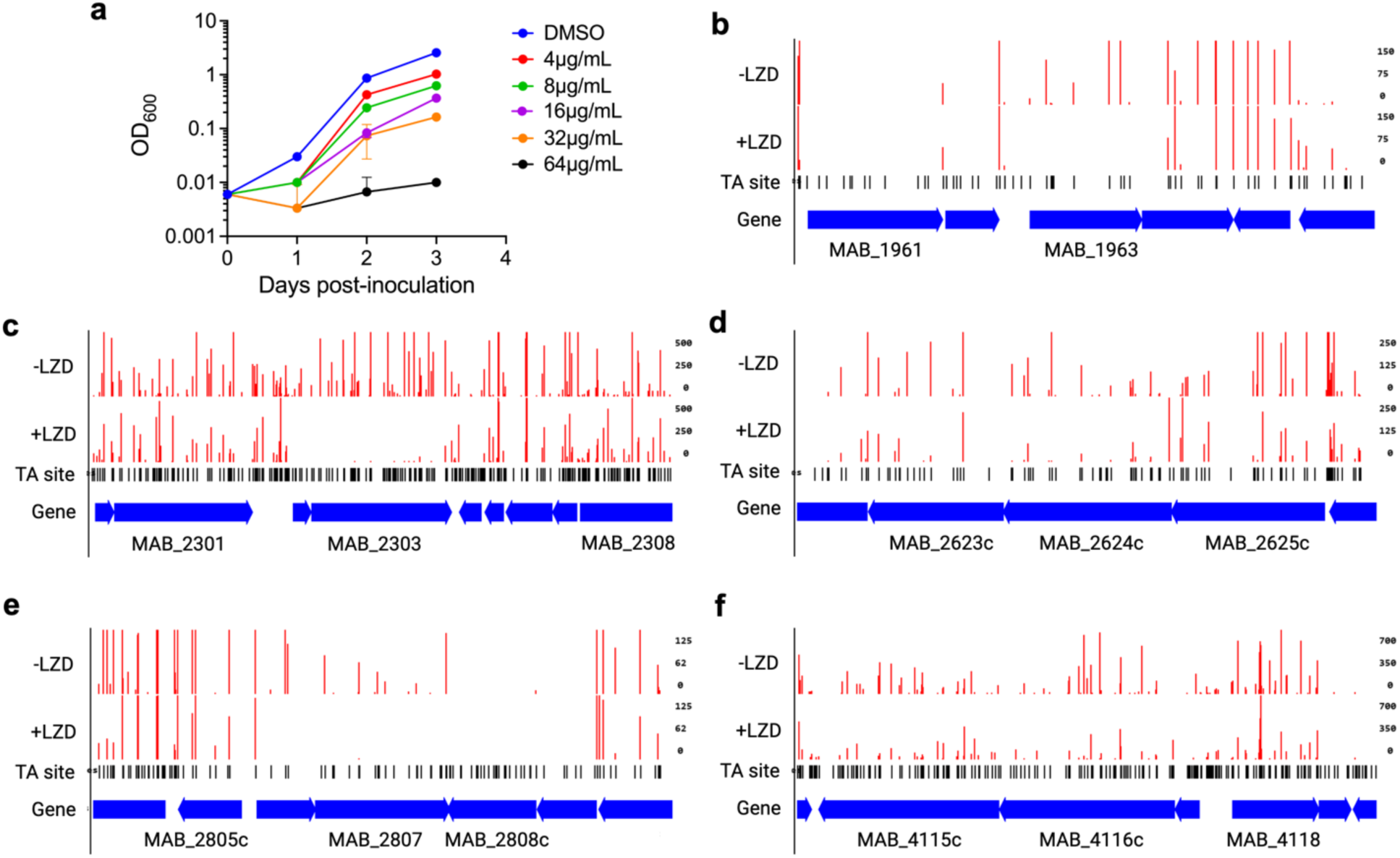
TnSeq screening in clinical isolate BWH-F reveals determinants of linezolid resistance. **a,** Relative growth as measured by optical density of *M. abscessus* clinical isolate BWH-F with specified concentrations of linezolid over time. Data are represented as individual values along with mean ± s.d. n=3 biological replicates. **b-f,** Transposon insertion counts for indicated genes in representative replicates of the -linezolid and +linezolid conditions. Insertion counts are normalized to the local maximum. DMSO = dimethylsulfoxide.

**Extended Data Fig. 3:**
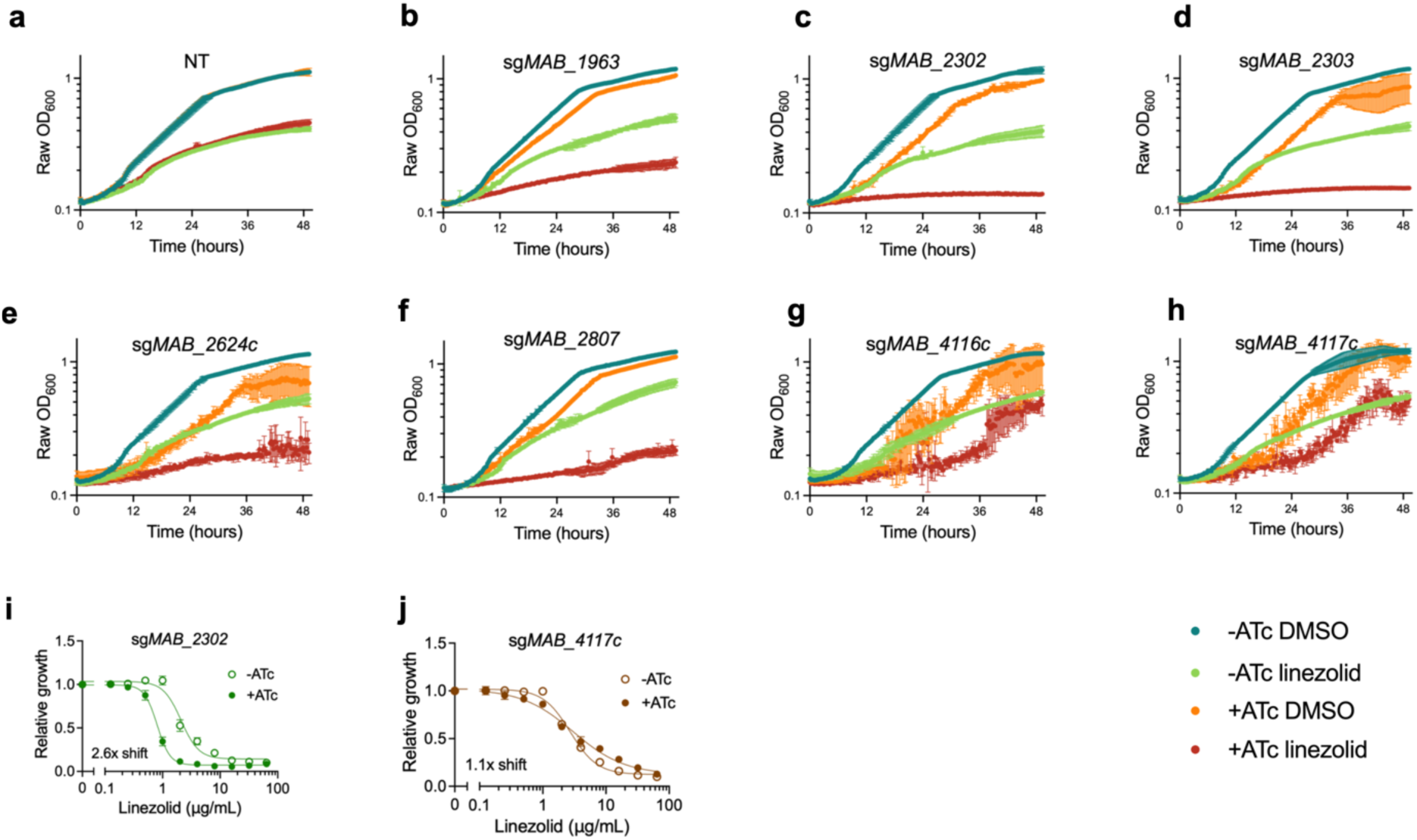
Knockdown down of membrane transporters increases sensitivity to linezolid. **a-h,** OD_600_ over time of pre-depleted *M. abscessus* ATCC19977 strains with sgRNAs targeting membrane transporter genes treated with 1μg/mL linezolid or vehicle along with ±ATc for 48 hours. **i-j,** Relative growth of *M. abscessus* ATCC19977 strains with sgRNAs targeting indicated membrane transporter genes as measured by a colorimetric dye treated with indicated concentrations of linezolid along with ±ATc for 48 hours. DMSO = dimethyl sulfoxide. ATc = anhydrotetracycline. NT = non-targeting sgRNA.

**Extended Data Fig. 4:**
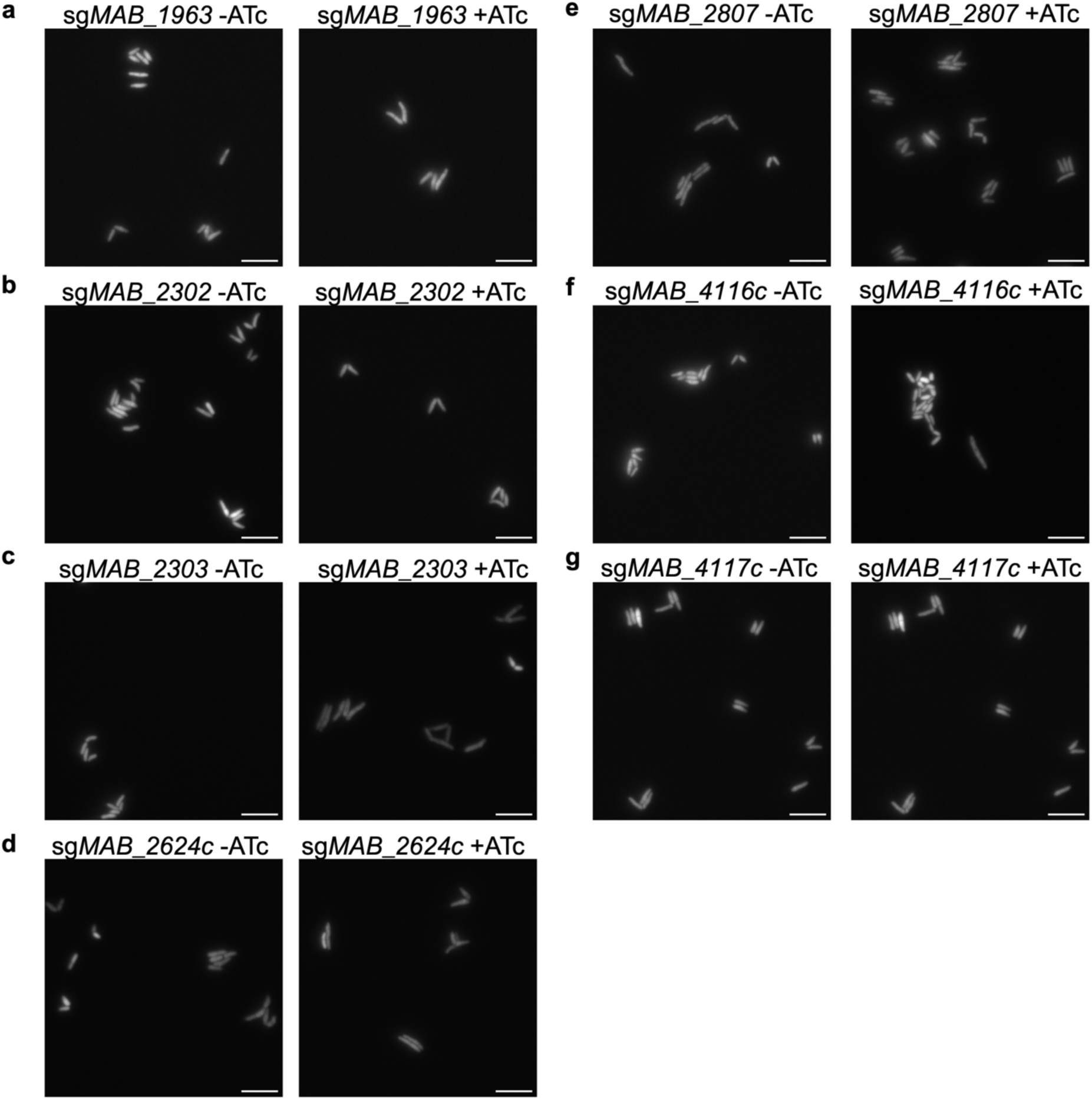
Membrane transporter mutants do not display gross morphological changes. **a-g,** Representative fixed cell widefield microscopy images of *M. abscessus* ATCC19977 strains with sgRNAs targeting membrane transporter genes in the presence or absence of ATc for 24 hours prior to fixation. Images were taken at 100x magnification. Scale bar = 5 µm.

**Extended Data Fig. 5:**
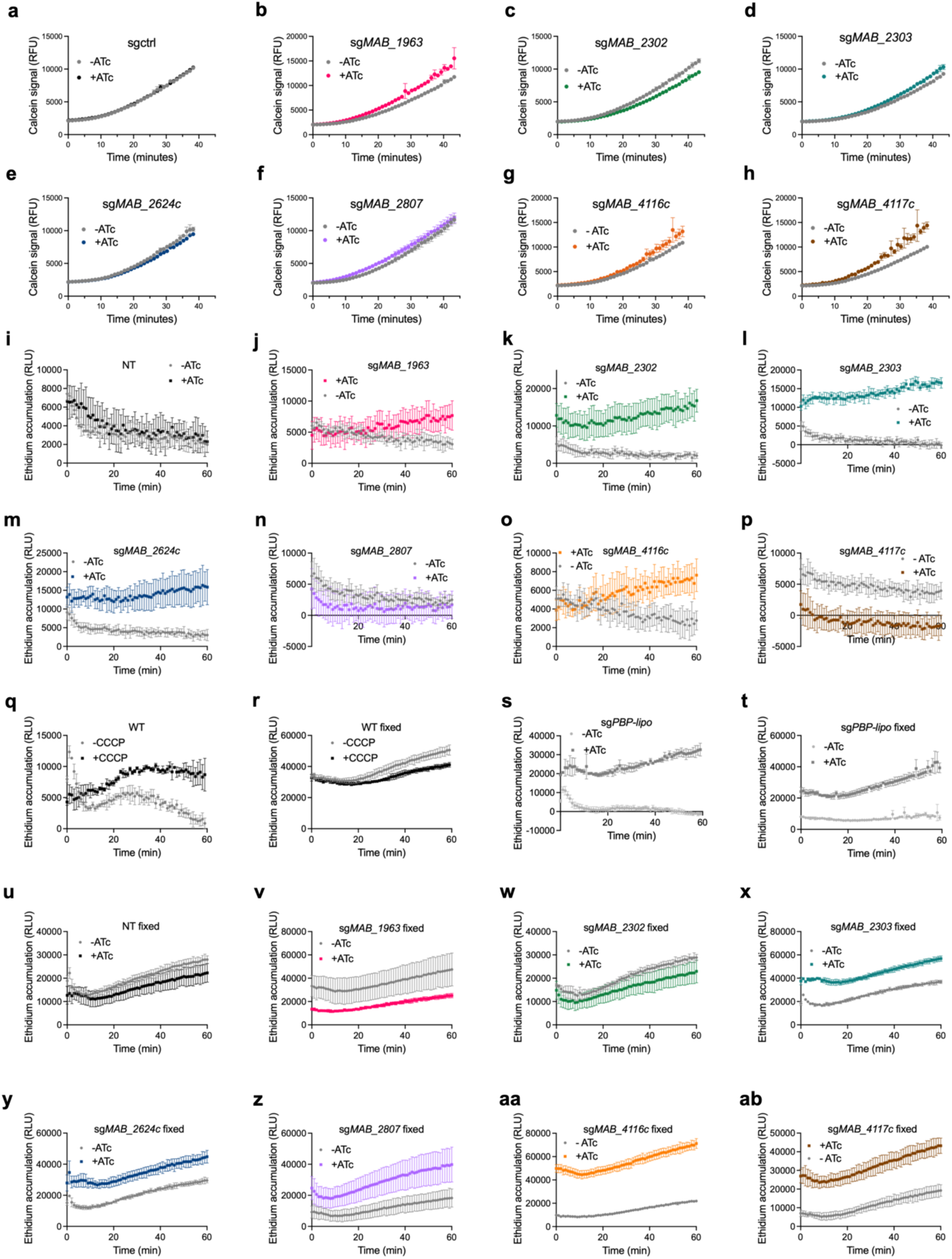
Accumulation of calcein and ethidium in membrane transporter knockdown strains. **a-h,** Calcein accumulation in *M. abscessus* ATCC19977 strains with sgRNAs targeting membrane transporter genes in the presence or absence of ATc for 24 hours prior to addition of calcein AM. Data are represented as individual values along with mean ± s.d. n=3 biological replicates. ATc = anhydrotetracycline. **i-p,** Ethidium accumulation as measured by fluorescence over time in live *M. abscessus* ATCC19977 strains with indicated sgRNAs targeting membrane transporter genes treated with 500ng mL^−1^ ATc for 24 hours prior to addition of ethidium bromide. **q-r,** Ethidium accumulation as measured by fluorescence over time in live and fixed wildtype *M. abscessus* ATCC19977 treated with 50μM CCCP. **s-ab,** Ethidium accumulation in (s) live or (t-ab) fixed *M. abscessus* ATCC19977 strains with indicated sgRNAs targeting membrane transporter genes pre-treated with 500ng mL^−1^ ATc for 24 hours. Data are represented as individual values along with mean ± s.d. n=3 biological replicates. ATc = anhydrotetracycline. WT = wildtype. NT = non-targeting sgRNA.

**Extended Data Fig. 6:**
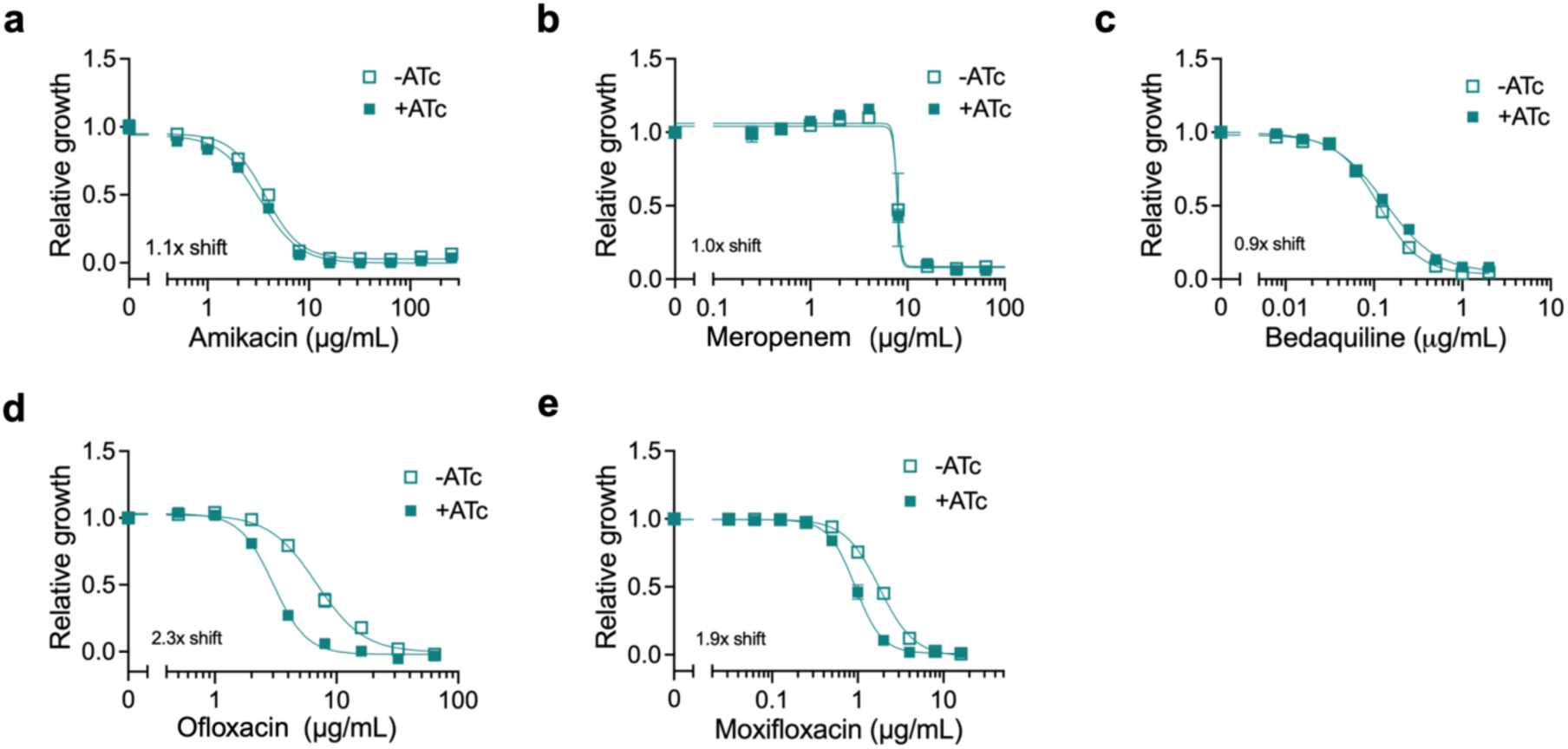
MAB_2303 only exhibits MIC shifts with chemically similar compounds. **a-e,** Relative growth of pre-depleted sg*MAB_2303 M. abscessus* strain as measured by reduction of a colorimetric dye after treatment with indicated concentrations of **(a)** amikacin, **(b)** meropenem, **(c)** bedaquiline, **(d)** ofloxacin, and **(e)** moxifloxacin in the presence or absence of 500 ng mL^−1^ ATc for 24 hours. Values normalized to vehicle only control per drug. Data are represented as individual values along with mean ± s.d. n=3 biological replicates. ATc = anhydrotetracycline.

**Extended Data Table 1:** Hits from TnSeq screen display moderate homology with other pathogenic mycobacterial species. Percent identity of amino acid sequences from *M. abscessus* with corresponding closest orthologs from *M. tuberculosis* H37Rv, *M. avium* 104, and *M. smegmatis* mc^2^155 as determined by alignment using Clustal 2.0.9 multiple sequence alignment.

| **Protein in *M. abscessus* ATC19977** | ***M. tuberculosis* H37Rv closest ortholog and % identity** | ***M. avium* 104 closest ortholog and % identity** | ***M. smegmatis* mc<sup>2</sup>155 closest ortholog and % identity** |
| --- | --- | --- | --- |
| MAB_1963 | Rv2198c, 54.42% | MAV_2293, 44.22% | MSMEG_4265, 56.38% |
| MAB_2303 | Rv0450c, 56.26% | MAV_2510, 55.42% | MSMEG_3496c, 54.55% |
| MAB_2624c | N/A | N/A | MSMEG_3249, 74.93% |
| MAB_2807 | Rv1410c, 62.82% | MAV_3369, 60.04% | MSMEG_3069, 65.67% |
| MAB_4116c | Rv0450c, 56.26% | MAV_3248, 67.37% | MSMEG_0381, 78.29% |

